# Elevated Temperatures Representing Heatwave Conditions Shift Active Nitrifying Communities and Their Viruses in Tidal Flats and Agricultural Soils

**DOI:** 10.1101/2025.10.17.682989

**Authors:** Baozhan Wang, Ping Gao, Ping Zhang, Yue Zheng, Xu Liu, Ning Ling, Jun Shan, Rongjiang Yao, Shuai Zhao, Zhiguo Zhang, Guibing Zhu, Man-Young Jung, Jianwen Zou, Xiaoyuan Yan, Sungeun Lee, Christina Hazard, Graeme W. Nicol, Jizhong Zhou, Yunfeng Yang, Yongguan Zhu, David A. Stahl, Michael Wagner, Yanzheng Gao, Jiandong Jiang, Wei Qin

## Abstract

Global heatwave intensification associated with climate change will impact the nitrogen cycle, yet its effect on specific nitrifier groups or their interactions with viruses remains unclear. Using ^13^CO_2_-DNA-based stable isotope probing (SIP) coupled with metagenomics, we show that elevated temperatures associated with heatwave conditions restructure active nitrifying communities and their viruses in Yangtze River estuary tidal flats and adjacent agricultural soils. In tidal flats, high temperatures shifted active ammonia-oxidizing archaea and bacteria (AOA and AOB), and nitrite-oxidizing bacteria (NOB) from marine to terrestrial ecotypes. In contrast, heatwave conditions stimulated terrestrial ecotypes of AOA but suppressed AOB in agricultural soils. ^13^C-labeled nitrifier-infecting viruses also showed temperature-driven shifts in activity, lifestyle, and auxiliary metabolic genes in concert with hosts. Notably, AOA viruses carried the plastocyanin gene, potentially augmenting host metabolism. These findings demonstrate heatwaves drive complex shifts in nitrifier communities and their interactions with viruses, impacting global nutrient cycling under climate extremes.

## Introduction

A heatwave is characterized as an extended period of unusually high temperatures, lasting from several days to months, with substantial effects on natural ecosystems, agricultural productivity, and human health ^1–5^. As global climate change accelerates, the frequency, intensity and duration of extreme heatwave events are increasing globally (IPCC, 2021). In many regions, heatwaves regularly reach temperatures exceeding 35-40°C with extremes reaching up to 45-50°C, and durations that, in severe cases, may extend over two months ^6–13^. Temperature is a critical driver of microbial metabolism, activity, and community structure, suggesting that heatwaves can drastically alter microbial community dynamics and interactions in terrestrial ecosystems ^14–19^. These changes can, in turn, profoundly influence biogeochemical cycles and ecosystem services.

Coastal wetlands, serving as dynamic transition zones between terrestrial and marine ecosystems, face dual pressures from both rapid climate change and extensive anthropogenic disturbances. They represent critical climate-sensitive regions on Earth, simultaneously receiving substantial inputs of reactive nitrogen from agricultural systems ^20, 21^. The intensive conversion of coastal wetlands into agricultural systems across many regions of the World further exacerbates significant repercussions on soil nutrient balance, making these vulnerable ecosystems even more fragile in the face of extreme climate conditions. However, the effects of heatwaves on nitrogen cycling − the nutrient having the most significant impact on local ecosystem functioning and global biogeochemical processes − remain poorly understood.

Ammonia and nitrite-oxidizing microorganisms (i.e., nitrifiers) are responsible for the oxidation of approximately 2,330 Tg of nitrogen (N) per year, constituting one of the largest nitrogen fluxes in the global nitrogen cycle budget ^22^. AOA and AOB convert ammonia (hereafter referred to as total ammonia, NH_3_ + NH_4_^+^) into nitrite. Nitrite is subsequently oxidized to nitrate by NOB, completing the two-step nitrification process. The recent identification of a novel group of nitrifiers, known as complete ammonia oxidizers (comammox, CMX), within the genus *Nitrospira*, demonstrates that some microbes are capable of oxidizing ammonia to nitrate in a single cell ^23, 24^. Although all nitrifiers generally depend on the energy generated from oxidation of ammonia and/or nitrite ^25^, some lineages exhibit metabolic versatility, such as the ability of utilizing urea, cyanate, and guanidine as ammonia sources ^26–28^. Some NOB can even grow on hydrogen instead of nitrite ^29, 30^. The distinct substrate preferences among different nitrifier lineages minimize direct substrate competition, enabling their coexistence in diverse environments ^26–28^. In addition to differences in N source utilization, major nitrifier lineages exhibit distinct thermal traits. For example, AOA and AOB generally show differences in temperature dependent nitrification activities in soils ^31–36^. However, the ecological implications of these thermal differences, particularly their responses to extreme heatwave events, remain largely unknown, despite their clear relevance to nitrifier community structure and the nitrification process in the context of ongoing climate change ^37^.

In addition to bottom-up controls of nitrification, one of the major top-down controls on bacterial and archaeal communities is viral infection. Viruses, the most abundant biological entities on Earth, play a pivotal role in regulating the community structure, metabolic activity, and evolutionary dynamics of bacterial and archaeal populations, thereby significantly influencing the biogeochemical cycles driven by prokaryotes ^16,38–42^. For instance, in marine ecosystems, an estimated 10^28^ viral infections occur daily, accounting for up to 50% of bacterial mortality ^43^, profoundly impacting prokaryotic ecology and biogeochemistry in the oceans. Viral infection facilitates the horizontal transfer of key metabolic and regulatory genes in nitrifiers. An auxiliary metabolic gene (AMG) encoding the archaeal AmoC subunit of ammonia monooxygenase, was first discovered in the global ocean virome ^44^, with virus-encoded *amoC* genes sometimes comprising up to half of total *amoC* gene copies in cellular fraction metagenomes ^45^. Given that previous physiological characterization has shown that AmoC likely maintains the integrity of functional AMO complex under stressed conditions ^46, 47^, viral AmoC may enhance the adaptive capacity of AOA hosts in oligotrophic marine environments. In contrast, viruses infecting soil AOA appear to lack *amoC* AMGs and contain other AOA-specific AMGs ^48, 49^. Furthermore, previous studies of pure cultures and environmental populations of nitrifier-infecting viruses, including those in lytic and lysogenic phases ^44, 45, 48–54^, suggests that viruses substantially affect the physiology and genomic dynamics of nitrifying microorganisms. However, our understanding on how viral infections regulate nitrifier ecology and nitrification activity remains limited. Even less is known about how viral infections interact with bottom-up controls, such as elevated temperatures, to influence the composition, activity and ecological function of nitrifying microorganisms.

Microcosm incubation are widely used to examine the effects of heatwave conditions on microbial communities ^55–57^. In this study, we employed ^13^C-DNA–based SIP with microcosms at 37 °C to investigate how heatwave conditions reshape active nitrifier communities, alter nitrification activity, and modulate virus–host interactions in two distinct coastal landscapes. We focused on natural upper tidal flats and adjacent agricultural soils that were converted from tidal flats in the 1960s at the estuary of the Yangtze River—the longest river in Asia—to explore how land-use change mediates nitrification and host–virus dynamics under heatwave conditions. We hypothesized that heatwaves would exert distinct effects on the structure of active nitrifying communities across land-use types and would strongly reconfigure virus–host interaction modes, revealing how bottom-up and top-down forces jointly shape nitrogen cycling under climate extremes.

## Material and Methods

### Site description and sample collection

The coastal upper tidal flat sampling site was located in Qidong City (30°14′N, 120°09′E), Jiangsu Province, China, within a typical subtropical monsoon climate zone. The region has a mean annual temperature of ∼15 °C and annual precipitation of 1037.1 mm. During midsummer (July–August), air temperatures average ∼28 °C, with 95% of values between 26–31 °C (Nantong City, China; https://berkeleyearth.org/ <u>temperature-location/31.35N-120.63E</u>). Summer heatwaves in the Yangtze River Delta can sustain temperatures >35 °C, with local extremes up to 45 °C, persisting for weeks to over two months ^7, 9, 12, 13^. The study site lies in upper tidal flats inundated only briefly during monthly spring tides (1–3 h), so soil temperatures at 0–15 cm closely track ambient air, differing by –0.7 ± 2.3 °C ^58^. Since the 1960s, ∼266,700 ha of tidal flats in Jiangsu Province have been reclaimed for agriculture. A paired sampling strategy was employed to collect three tidal flat and agricultural soil samples at a depth of 0-15 cm along the northern branch of the Yangtze River estuary from west to east in summer 2018 (Supplementary Fig. S1). At each site, five soil cores were randomly collected within a 10 × 10 m plot, transported on dry ice, freeze-dried, sieved (2.0 mm), homogenized, and combined into a composite sample, which was stored at –20 °C until analysis.

### 13CO2-DNA-SIP microcosms

Soil physicochemical properties were determined as described previously ^59–62^. Nitrification potential rates were assayed at 5 °C intervals from 5–40 °C following established protocols^63^. DNA-SIP microcosms were set up to identify active nitrifiers ^59, 60^. Triplicate 120-ml serum bottles each contained 12.0 g sieved fresh soil (≈10.0 g dry weight), amended with 50 µg NH_4_Cl-N g_−1_ dry soil, approximating *in situ* concentrations. Bottles were sealed and injected with ^13^CO_2_ (99 at% C; Sigma-Aldrich) or ¹²CO_2_ (generated from Na_2_CO₃ acidification) for a final concentration of 5% (v/v), and incubated at 28 and 37 °C in the dark for 8 weeks at 60% water-holding capacity ^64, 65^. Headspaces were flushed with synthetic air (20% O_2_, 80% N_2_) and replenished with CO_2_ every 4 days. Ammonium, nitrite, and nitrate concentrations were measured weekly ^60^, and NH_4_Cl-N was re-amended when concentrations fell below 6 µg g_−1_ dry soil.

After incubation, DNA was extracted from 0.5 g soil using the FastDNA Spin Kit for Soil (MP Biomedicals). Approximately 5 µg DNA was fractionated by CsCl ultracentrifugation (initial density 1.725 g ml_−1_; 177,000 g, 44 h, 20 °C; VTi65.2 rotor, Beckman Coulter). Fourteen fractions (∼360 µl each) were collected by syringe pump (New Era Pump Systems), buoyant densities determined with an AR200 refractometer (Reichert), and DNA recovered by PEG 6000 precipitation before dissolution in 30 µl TE buffer ^66^. The abundance of *amoA* genes from AOA, AOB, and CMX in all fractions, as well as in pooled heavy DNA fractions (1.733–1.745 g ml_−1_), was quantified by qPCR using a CFX96 system (Bio-Rad). Further details are provided in Supplementary Methods.

### Metagenomic sequencing and assembly

Metagenomic sequencing was performed on total DNA from original soil samples and pooled heavy DNA fractions (1.733–1.745 g ml_−1_) ^59, 60^ from both ^13^CO_2_-labeled and ¹²CO_2_-control treatments using the Illumina NovaSeq platform (Shanghai Majorbio Bio-Pharm Biotechnology Co., Ltd., Shanghai, China). Assembly, binning, and annotation were conducted as described in Supplementary Methods. In total, 28 AOA, 21 AOB, 3 CMX, and 15 NOB MAGs of medium (≥50% completeness, ≤10% contamination) to high quality (≥90% completeness, ≤5% contamination) were recovered from *in situ* (total original soil DNA) and ^13^C-DNA metagenomes. Phylogenomic and comparative genomic analyses are detailed in Supplementary Methods.

### Construction of in-house AmoA/NxrB database and community structure analysis

In addition to *amoA* and *nxrB* genes recovered from this study’s metagenomes, homologs were compiled from public nitrifier genomes (1,698 AOA, 528 AOB, 903 NOB/comammox) in NCBI RefSeq, GTDB (April 2024), KEGG, and previous datasets ^23, 67, 68^. Redundant sequences were dereplicated with CD-HIT ^69^ at 100% identity to generate a non-redundant amino acid dataset, which was aligned with MAFFT (version 7.407) ^70^, trimmed with Gblocks (version 0.91b) ^71^, and used to construct phylogenies in IQ-TREE (version 2.2.0) ^72^ with automatic model selection and 1,000 ultrafast bootstraps, ensuring consistency with established nitrifier lineages ^23, 67, 68, 73–76^. Community structure in ^13^C-labeled SIP microcosms was assessed by mapping quality-filtered reads to this database with DIAMOND (version 2.0.2) ^77^ (≥80% identity, ≥100 bp coverage, e-value ≤1×10_−1_⁰), assigning them to lineages based on phylogenetic placement, and normalizing by sequencing depth.

### Identification, taxonomy and lifestyle of the ^13^C-labeled vOTUs

Putative ^13^C-labeled viral contigs ≥5 kb and ≥10 kb were identified with VirSorter2 v2.2.3 ^78^ at a loose cutoff of 0.5 for maximal sensitivity ^79, 80^. Sequences were quality-checked with CheckV v0.9.1 ^81^, requiring at least one viral hallmark gene (e.g., capsid, portal, terminase, integrase) or enrichment in unannotated genes ^49^, and potential host-derived regions were trimmed. Viral contigs sharing ≥95% nucleotide identity across ≥85% of the shorter contig were clustered into vOTUs using Cluster_genomes_5.1.pl ^82^, with the longest contig in each cluster selected as representative. Taxonomic assignment was performed with PhaGCN2 v2.1 ^83^ following ICTV guidelines, and viral lifestyle (lytic vs. lysogenic) was predicted with DeePhage v1.0 ^79, 84, 85^ at a 0.5 cutoff.

### Virus-host linkage, gene-sharing network, and 3D structure prediction of viral plastocyanin

“*Ex situ*” hosts of ^13^C-labeled viral contigs were inferred from public nitrifier genomes (1,698 AOA, 528 AOB, 903 NOB; NCBI and GTDB, Sept 2024), and “*in situ*” hosts from the 67 nitrifier MAGs recovered in this study ^86^. Host prediction combined three in silico approaches: (i) CRISPR matching—CRISPR spacers (≥25 bp) were extracted with CRT v1.1 ^87^, dereplicated at 100% identity (CD-HIT v4.8.1 ^69^), and queried against viral contigs (≥100% identity, ≤1 mismatch, e-value ≤10⁻⁵); (ii) homology-based inference—viral proteins were compared against the NCBI nr database with DIAMOND BLASTp (e-value <10⁻⁵), and a viral host was assigned to nitrifiers when AOA, AOB, or NOB genomes accounted for the largest proportion of ORF “best-hit” ^88^; and (iii) tRNA matching—host and viral tRNAs were predicted using tRNAscan-SE v1.3.1 ^89^, with viral tRNAs linked to nitrifiers only if uniquely present in nitrifiers and identical (or with ≤1 mismatch) to host tRNAs. Viral contigs supported by at least one method were classified as nitrifier-infecting. A gene-sharing network of nitrifier-infecting viruses was constructed using vConTACT2. The 3D structure of viral plastocyanin was predicted with I-TASSER-MTD (https://seq2fun.dcmb.med.umich.edu/I-TASSER/) and visualized and aligned with reference proteins using PyMOL v3.12 ^90^.

### Statistical analysis

All analyses were conducted in R v4.2.1. Independent-sample *t*-tests compared the relative abundances of nitrifier-infecting viruses versus hosts and the frequencies of function-specific AMGs in ^13^C-labeled metagenomes across temperature treatments. Effect sizes were calculated as Cohen’s *d* ^91^ (mean difference/pooled SD), with AMG frequency defined as each AMG’s RPKM relative to the total RPKM of all AMGs.

Effect sizes were classified as large (|*d*| > 0.8), medium (0.5 < |*d*| ≤ 0.8), small (0.2 < |*d*| ≤ 0.5), or negligible (|*d*| ≤ 0.2).

## Results

### Temperature-driven shifts in ^13^C-labeled active nitrifiers in tidal flats and agricultural soils

At the study site, midsummer (July–August) air temperatures average ∼28 °C, with 95% of values between 26–31 °C (Nantong City, China, https://berkeleyearth.org/temperature-location/31.35N-120.63E), while heatwaves frequently elevate temperatures to 35–40 °C for weeks to two months ^7, 9, 12, 13^. The sampling area lies in upper tidal flats that are inundated only briefly during monthly spring tides (1–3 hours per month), so soil temperatures (0-15 cm) closely track ambient air, differing by –0.7 ± 2.3 °C ^58^. Nitrification potential assays revealed contrasting thermal optima: tidal flat communities peaked at 25–30 °C, whereas adjacent agricultural soils—converted from tidal flats in the 1960s—showed maximal activity at 35–40 °C (Supplementary Fig. S1). These patterns are to a large degree consistent with cultured representatives, in which marine ecotypes typically display optima of ∼20–32 °C ^92–96^ and terrestrial ecotypes of ∼35–42 °C ^35, 36, 97–100^. Microcosm incubations are usually used to examine microbial responses to heatwave conditions ^55–57^, therefore guided by these climatic and physiological observations, we conducted ^13^C-DNA SIP microcosm incubations at 37 °C to mimic prolonged heatwave conditions and at 28 °C to represent average midsummer temperatures, enabling us to disentangle the respective effects of land-use history and elevated temperature warming on nitrifier community structure and virus–host interactions.

At 28°C, AOB outnumbered AOA at both tidal flat and adjacent agricultural sites, with CMX detected only in agricultural soils and nearly absent in tidal flats. After eight weeks of incubation, nitrification activity at both temperatures aligned well with the optimal temperatures identified for each habitat, with tidal flats and agricultural soils accumulating more nitrite/nitrate (NO_x_) at 28°C and 37°C, respectively (Fig. 1A and Supplementary Fig. S1). Subsequent extraction of total DNA from the ^13^C-labeled and ^12^C-control DNA-SIP microcosms, followed by ultracentrifugation and qPCR quantification of the *amoA* gene (encoding the alpha submit of the AMO), revealed differential labeling of bacterial and archaeal ammonia oxidizers in ^13^C-labeled heavy DNA fractions at 28°C and 37°C (Fig. 1B and Supplementary Fig. S2). For instance, in agricultural soils, the abundance of AOB in the ^13^C-DNA was decreased at 37°C compared to 28°C (*P* < 0.01 for Agr-1), and the ^13^C-labeled CMX abundance was also reduced in the three samples (*P* < 0.001) at the elevated temperature to very low levels (less than 1.0 × 10^4^ copies g^-1^ dry weigh soil) (Fig. 1B). In contrast, the abundance of ^13^C-labeled AOA more than tripled, or even increased more than fivefold, at 37°C relative to 28°C, becoming the dominant and active ammonia oxidizers under sustained elevated temperature (Fig. 1B). These findings indicated that AOA activity was stimulated under sustained high temperature conditions in agricultural soils, suggesting their greater thermal adaptability compared to bacterial ammonia oxidizers. In contrast, in tidal flats, sustained heat at 37°C had overall negative effects on all three major ammonia oxidizer lineages. CMX represented only a minor ammonia-oxidizing microbes in tidal flats, with ^13^C-labeled comammox abundance falling below the detection limit (< 1.0 × 10^3^ copies g^-1^ dry weigh soil) at 37°C. For AOB, there was a notable decrease in ^13^C-labeled abundance at 37°C compared to 28°C, with a reduction ranging from 43.8% to 47.1%. While also showing a decreasing trend with increased temperature, tidal flat AOA exhibited relatively higher tolerance to sustained heat, with two of the three incubation groups (Tid-1 and Tid-3) showing no significant difference in ^13^C-labeled abundance between 28°C and 37°C (*P* > 0.05) (Fig. 1B). Within each major ammonia oxidizer lineage, we observed a major shift in active subpopulations of AOA and AOB as temperature increased (see below).

**Figure 1.**
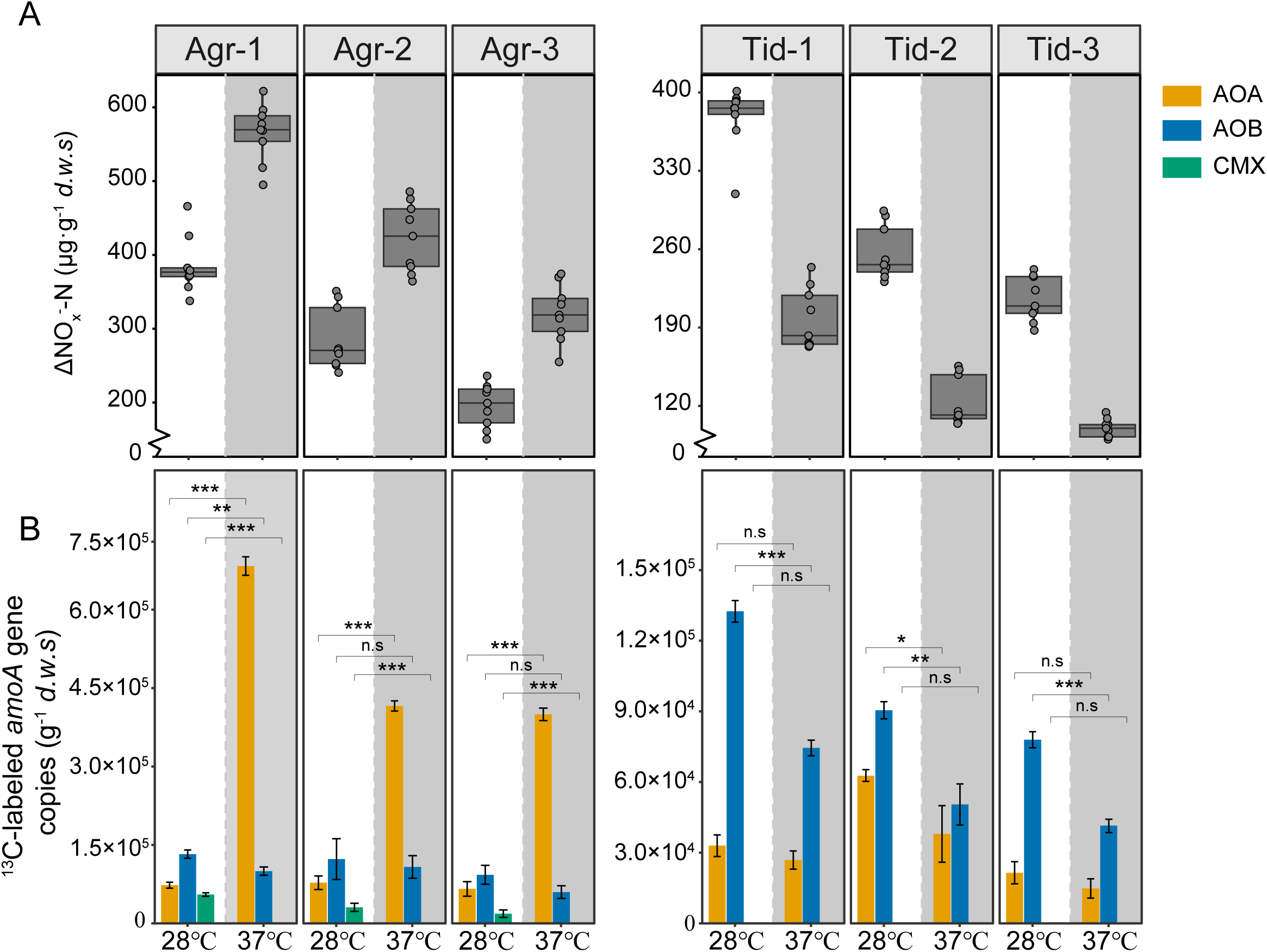
Nitrification activities and ^13^CO_2_-labeling of ammonia oxidizers in soil microcosms over 8 weeks at different temperatures. **(A)** Changes in soil NO ^-^ (the combination of nitrite and nitrate) concentrations were analyzed to assess nitrification activity in soil microcosms at different temperatures. Each box represents the middle 50th percentile of the data set and is derived using the lower and upper quartile values, with whiskers extending to the 5th–9th percentiles. The median value is indicated by the center line in each box. Outliers are depicted as circles. **(B)** The *amoA* gene copy numbers of AOA (yellow bars), AOB (blue bars) and comammox (green bars) in the ^13^C-labeled DNA were determined using real-time quantitative PCR. Temperature treatments are indicated along the x-axis, with the “37°C” fractions highlighted by shaded rectangles in grey. All treatments were conducted in triplicate microcosms, and error bars represent the standard deviation of the triplicate experiments.

### Community shift of ^13^C-labeled nitrifiers at high temperature

Metagenomic analysis of the ^13^C-DNA from DNA-SIP further substantiated distinct community shifts in responses to temperature differences between active nitrifiers in agricultural soils and tidal flats (Fig. 2). The relative abundance of major ammonia-oxidizing microbial lineages estimated from metagenomic analysis was well aligned with qPCR quantification, supporting a temperature-driven transition in agricultural soils from AOB dominance at 28 °C to AOA dominance at 37 °C (Fig. 2A).

**Figure 2.**
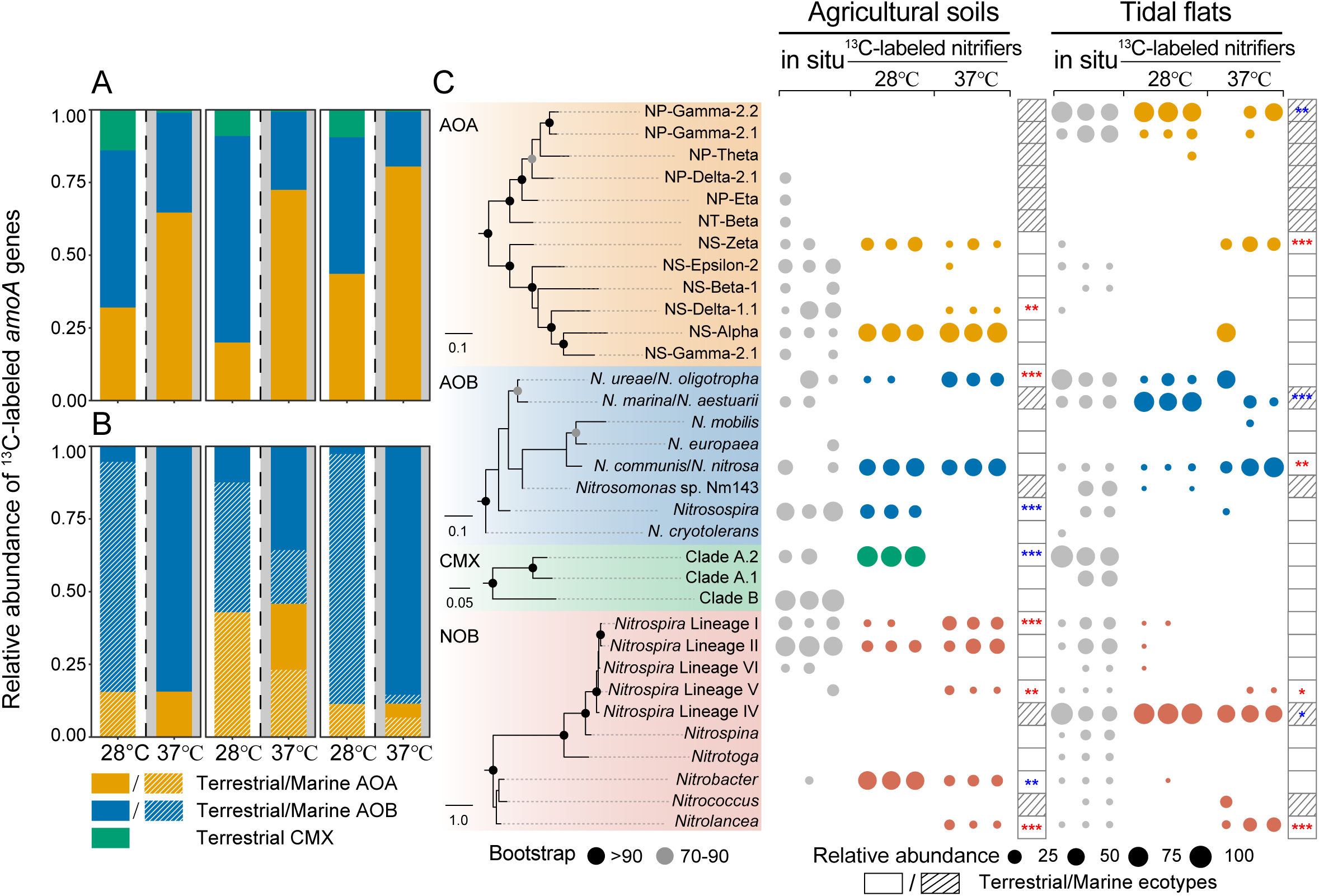
Population dynamics of the ^13^C-labeled functional nitrifiers at different temperatures. **(A-B)** Relative abundance of *amoA* genes from AOA, AOB, and comammox in the ^13^C-labeled DNA from agricultural soils (**A**) and tidal flats (**B**) at different temperatures, as determined by a metagenomic read-recruiting approach. The designations alongside the x-axis represent different temperatures, with the “37℃” fractions highlighted by shaded rectangles in grey. Striped bars represent the relative abundance of typical marine ecotypes, whereas non-striped bars represent typical terrestrial counterparts. For comammox no marine ecotypes are described**. (C)** Phylogenetic trees of AOA, AOB, comammox *amoA* genes, and NOB *nxrB* genes in the ^13^C-labeled DNA metagenomes are presented on the left of panel c. These trees were conducted using IQ-TREE (version 2.1.2) ^84^ with the best-fit model selection. The major lineages within the trees have been collapsed for clarity. NP represents *Nitrosopumilales*, and NS represents *Nitrososphaerales* in the AOA phylogenetic tree. A bubble plot shows the composition of nitrifiers in ^13^C-labeled DNA and *in situ* DNA (total original soil DNA, grey) on the right of panel c, using a metagenomic read-recruiting approach for the *amoA* or *nxrB* genes. The number of asterisks within the rectangles signifies the level of statistical significance (\**p* < 0.05, \*\**p* < 0.01, \*\*\**p* < 0.001), with colors indicating an increase (red) or decrease (blue) in relative abundance as temperatures rise. Rectangles with stripes on the right represent typical marine types, while those without stripes denote typical terrestrial types.

Competitive fragment recruitment of ^13^C-labeled metagenomes revealed that active nitrifying groups, including AOA, AOB, CMX, and canonical NOB, exclusively belonged to terrestrial ecotypes at both temperatures in agricultural settings (Fig. 2C). For instance, all ^13^C-labeled AOA belonged to the soil order *Nitrososphaerales*, predominantly composed of NS-Zeta and NS-Alpha sublineages in the labeling experiment, irrespective of temperature (Fig. 2C). Among AOB, *N. communis*/*N. nitrosa* was the most abundant sublineage at both temperatures, while the *Nitrosospira* sublinaege was dominant only at 28°C but was undetected at 37°C. In contrast, the relative abundance of *N. ureae*/*N. oligotropha* increased significantly (*P* < 0.001) at elevated temperature (Fig. 2C). For NOB, ^13^C-labeled *nxrB* genes primarily associated with *Nitrobacter* and *Nitrospira* lineage II at 28°C. However, at 37°C, the NOB community became more diversified, with a significant increase in the relative abundance of *Nitrospira* lineages I and V, as well as *Nitrolancea* (*P* < 0.001) (Fig. 2C). The ^13^C-labeled CMX, detected only at 28°C, exclusively affiliated with Clade A.2 (Fig. 2C), while clade B members showed no labeling at both temperatures.

Unlike the converted agricultural fields, tidal flats are transitional zones in estuarine ecosystems, harboring a mix of marine and terrestrial nitrifier populations. These habitats are characterized by high salinity, with electrical conductivity ranging from 493 to 1,974 μs/cm (Supplementary Fig. S1). The sampling sites are only briefly inundated by seawater during the monthly spring tide (1–3 hours per month), so the surface temperature of the tidal flats for most of the month is nearly identical to the ambient air temperature, differing by only –0.7 ± 2.3 °C ^58^. We observed that sustained high temperature in tidal flats led to a significant shift in functional nitrifier communities, with active terrestrial ecotypes emerging at 37°C but absent at 28°C (Fig. 2B and C). Specifically, at 28°C, all active AOA in the tidal flats belonged to the typical marine groups, NP-Gamma and NP-Theta (Fig. 2C). However, incubation at 37°C stimulated the activity and increased the relative abundance of the terrestrial group NS-Zeta and NS-Alpha in tidal flats, which accounted for 31-100% of the total ^13^C-labeled AOA populations in the three samples (Fig. 2C). Likewise, when comparing incubations at 37°C to 28°C, the most active and dominant AOB ecotypes shifted from typical marine ecotypes *N. marina*/*N. aestuarii* to terrestrial ecotypes *N. communis*/*N. nitrosa*. (Fig. 2C). In addition, at both incubation temperatures, the ^13^C-labeled active NOB predominantly belonged to *Nitrospira* lineage IV, which is widely detected in marine environments (Fig. 2C), whereas the ^13^C-labeled *Nitrolancea*, a typical terrestrial NOB sublineage, was detected only at 37°C (Fig. 2C).

A total of 67 medium to high-quality metagenome-assembled genomes (MAGs) of nitrifiers were recovered from both ^13^C-labeld DNA and unlabeled soil metagenomes, including 28 AOA, 21 AOB, 3 CMX, and 15 canonical NOB MAGs (Supplementary Fig. S3-6, Supplementary Tables S1-2). Comparative genomic analysis revealed that temperature and salinity are major selective forces driving shifts in active nitrifying members in both agricultural and tidal flat ecosystems. For instance, nearly all of the ^13^C-labeled nitrifiers from the two environments, regardless the incubation temperature, shared common genes associated with salt tolerance. These included *cpa2* (Na^+^:H^+^ antiporter-2) for AOA (Supplementary Fig. S7 and Supplementary Table S3), *nqrABCDEF* (Na^+^-transporting NADH:ubiquinone oxidoreductase), and *phaACDEFG* (multicomponent K^+^:H^+^ antiporter) for AOB (Supplementary Fig. S8 and Supplementary Table S3), and *cpa2*, *yggT* (YggT family Na^+^/H^+^ antiporter), and *opuE* (solute:Na^+^ symporter, SSS family) for NOB (Supplementary Fig. S9 and Supplementary Table S3). Beyond these shared genes, marine ecotypes from tidal flats exhibited unique osmoregulatory genes. For instance, *Nitrosopumilales* contained *ectABCD* (genes involved in ectoine biosynthesis) and *nhaP2* (K^+^/H^+^ antiporter) (Supplementary Fig. S7), while *Nitrosococcus marina* and *Nitrosococcus aestuarii* featured *betL* (glycine betaine transporter) and *betT* (choline/glycine/proline betaine transport) genes (Supplementary Fig. S8). Additionally, *Nitrospira* lineage IV possessed *opuD/betL* (glycine betaine transporter) (Supplementary Fig. S9), all of which were absent in their terrestrial counterparts (Supplementary Fig. S7-10).

Furthermore, at 37°C, AOA and NOB terrestrial lineages in both agricultural soils and tidal flats displayed unique thermotolerance-related genes. For example, *Nitrososphaerales* AOA contained *mngA* (mannosyl-3-phosphoglycerate synthase) ^101, 102^ and *htpX* (heat shock protein) (Supplementary Fig. S7), *Nitrospira* lineages (excluding lineage IV) harbored *mngA* ^101, 102^ (Supplementary Fig. S9), and *Nitrolancea* NOB carried *DPH4* (DNAJ heat shock N-terminal domain-containing protein) (Supplementary Fig. S9). These genes were absent in counterparts whose activities were significantly reduced at 37°C compared to 28°C (Supplementary Fig. S7 and S9).

### Temperature-driven shifts in viral communities and virus-nitrifier interactions in agricultural soils and tidal flats

A total of 584 viral contigs ≥5 kbp in length, including 198 contigs ≥10 kbp, were recovered from the ^13^C-labeled metagenomes following an 8-week DNA-SIP microcosm incubation, and to minimize potential biases from short contigs, the following analyses were cross-validated with the ≥10 kbp subset. Clustering at 95% sequence identity and 85% coverage yielded 484 viral operational taxonomic units (vOTUs; Supplementary Table S4), of which 28.8% (158 of 584) were annotated to families under the ICTV classification, while the remainder were unclassified (Fig. 3A). The ^13^C-labeled viral communities exhibited clear temperature-dependent shifts: in agricultural soils, Peduoviridae and Mesyanzhinovviridae dominated at 28 °C, whereas Vilmaviridae was strongly enriched at 37 °C; in tidal flats, Mesyanzhinovviridae dominated at 28 °C, but Peduoviridae and Herelleviridae became more abundant at 37 °C. Host-linkage predictions using CRISPR spacer matches, tRNA gene similarity, and homologous gene searches identified at least 39, 63, and 60 contigs as viruses infecting AOA, AOB, and NOB, respectively (Fig. 3B; Supplementary Table S4). Collectively, nitrifier-infecting viruses accounted for ∼50% of the abundance of all ^13^C-labeled viral communities across both ecosystems and temperatures, and these patterns were further validated with the ≥10 kbp dataset (Supplementary Fig. S11 A and B). Intriguingly, four viral contigs linked to predatory Myxococcales and one to Bdellovibrionales were also detected, suggesting trophic interactions in which predators consumed ^13^C-labeled nitrifiers or heterotrophs that incorporated the label through cross-feeding, thereby revealing a complex microbial food web ultimately fueled by autotrophic nitrification.

**Figure 3.**
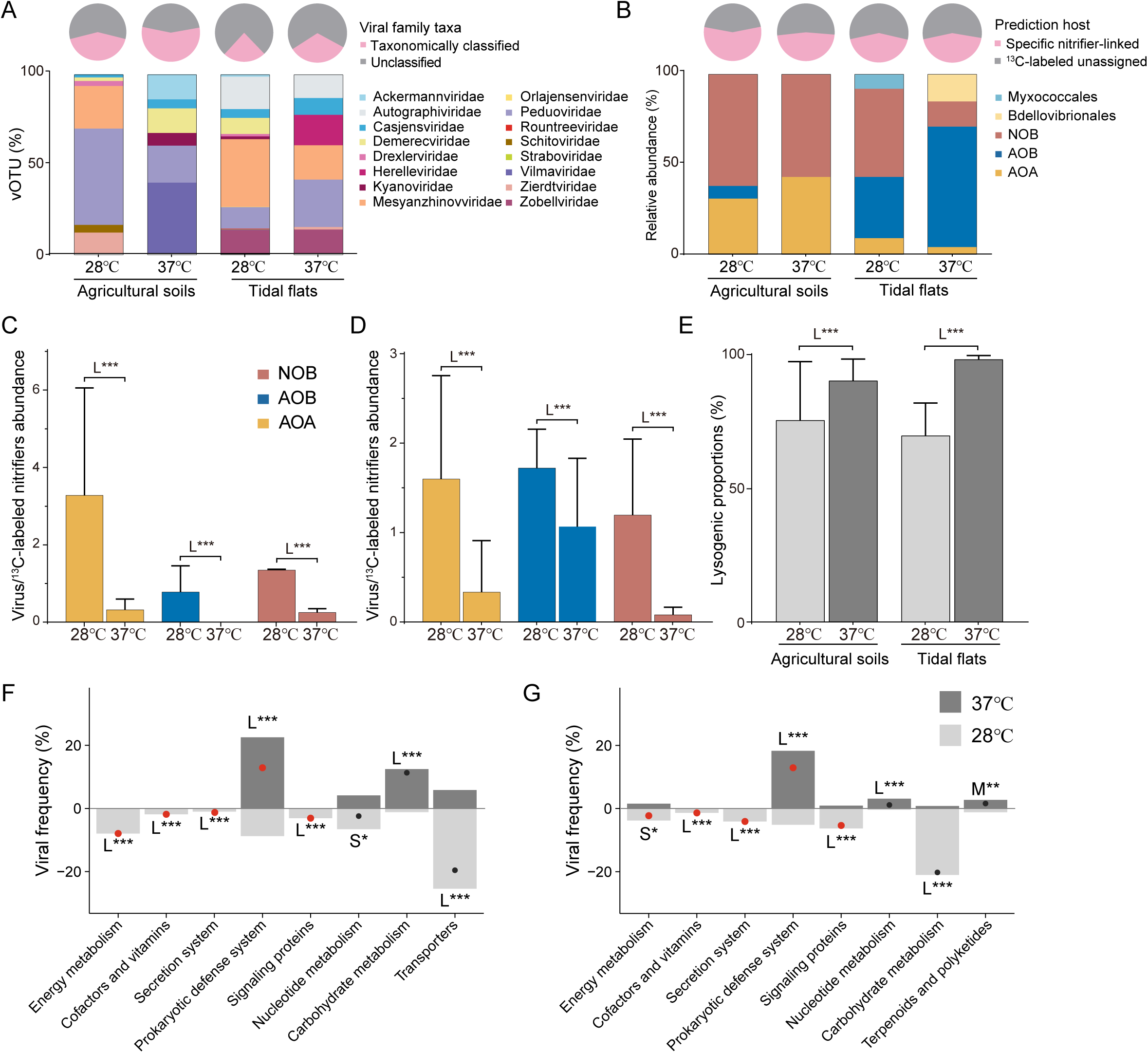
Characterization of the ^13^C-labeled viruses under varying temperatures in soil microcosms. **(A)** Taxonomic compositions of the ^13^C-labeled viruses in the agricultural soils and tidal flats at different temperatures. The lower bar charts illustrate the abundance of vOTUs with predicated taxonomy at the family level, based on the ICTV (International Committee Taxonomy of Viruses) classification, relative to total ^13^C-labeled viruses. The upper pie charts depict the relative abundance of summed taxonomically classified viruses within the total ^13^C-labled virus population. **(B)** Relative abundances of viral contigs predicted to infect AOA, AOB, canonical NOB and predator bacterial host populations, categorized as specific nitrifiers-linked viruses within the total ^13^C-enriched viral communities in this study. The bottom bar charts display the relative abundance of viral contigs from viruses infecting AOA, AOB, canonical NOB, or predator bacterial hosts relative to the total nitrifiers-linked viruses. **(C and D)** Relative abundance ratios of the nitrifier-infecting viruses to their corresponding hosts (AOA, AOB or NOB) in the ^13^C-labeled metagenomes of agricultural soils **(C)** and tidal flats **(D)** at different temperatures, calculated based on the accumulated coverage depths from read mapping to viral contigs to their corresponding host marker genes, such as the *amoA* genes from AOA and AOB, and *nxrB* gene from NOB, respectively. **(E)** The changes of lysogenic proportions of the ^13^C-labeled viruses as the temperature increased in agricultural soils and tidal flats ecosystems. **(F and G)** Frequency of viruses containing specific AMGs at 28℃ and 37℃. Red or black dots represent changes (37℃ minus 28℃) in agricultural soils (F) and tidal flats (**G**). In panels **(C-G)**, temperature effect sizes are classified as large (L***, |d| > 0.8), medium (M**, 0.5 < |d| ≤ 0.8), and small (S*, 0.2 < |d| ≤ 0.5) using Cohen’s d, with mean values and standard deviations indicated by error bars.

Under high-temperature conditions, virus–nitrifier interactions showed coordinated shifts in infection intensity, lifestyle strategies, and functional repertoires, indicating temperature-driven restructuring of virus–host dynamics. Virus-to-host abundance ratios were markedly higher at 28 °C than at 37 °C in both agricultural soils and tidal flats (Cohen’s d > 0.8; Fig. 3C and D), suggesting stronger virus–nitrifier interactions at 28 °C, whereas warming may reduce infection or favor more stable associations.

Elevated temperature (37 °C) also increased the proportion of lysogenic ^13^C-labeled viruses in both ecosystems (Cohen’s d > 0.8; Fig. 3E), although this trend was not significant in agricultural soils when restricting analyses to ≥10 kbp contigs, likely due to the smaller dataset (Supplementary Fig. S11C–E). Functional analyses further revealed that ^13^C-labeled viruses at 28 °C encoded a higher proportion of AMGs linked to energy metabolism (Cohen’s d = 1.63 and 0.47 for agricultural soils and tidal flats, respectively), cofactors and vitamins (0.82 and 0.97), secretion systems (1.63 and 1.04), and signaling proteins (67.45 and 0.99) (Fig. 3F, G; Supplementary Table S5), categories typically associated with active replication and lytic infections. In contrast, host defense functions related genes, also classified as borderline AMGs such as site-specific DNA methyltransferases and type II toxin–antitoxin systems, were enriched at 37 °C (Cohen’s d = –1.93 and –5.90; Fig. 3F, G; Supplementary Table S5). These findings closely matched those obtained with the ≥10 kbp dataset, underscoring the robustness of the observed trends (Supplementary Figure S11F and G). Collectively, these results suggest that under warming, viruses preferentially encode functions that maintain lysogeny, protect hosts from superinfection, and promote virus–host stability ^103^.

In addition to the functional viral contigs identified in this study, we performed a gene-sharing network analysis that incorporated a comprehensive reference dataset, including 324 previously reported nitrifier viruses, 2469 proviruses identified in 3,121 nitrifier genomes from the NCBI and JGI databases, and 2411 viral contigs predicated to infect nitrifiers from the IMG-VR v4 database (Supplementary Fig. S12A). Our analysis revealed that 27.7% of the viral contigs recovered in this study were integrated into the resulting network (Supplementary Fig. S12A). As expected, viruses infecting the same group of nitrifiers tended to cluster together (Supplementary Fig. S12A). Intriguingly, the network analysis also indicated that viruses associated with phylogenetically distinct nitrifying groups co-occurred within the same clusters, suggesting potential cross-infection (Supplementary Fig. S12A).

Notably, one viral contig (300022743|Ga0228699_1000093) within such a cluster was identified as infecting both NOB and AOB, which was supported by a spacer-to-protospacer match with NOB, and by sharing the most abundant ‘best hit’ homologues genes with AOB (Supplementary Fig. S12B).

### Viral plastocyanin genes associated with copper-dependent energy metabolism of AOA

One AOA-infecting viral contig, designated as contig vContig_022 (7.1 kbp), was retrieved from the ^13^C-labeled metagenome of agricultural soil microcosms incubated at 28°C. It contained a homolog of the AOA-specific plastocyanin (*pcy*) gene, which encodes one type of a single-domain cupredoxin ^104^, along with virus-specific genes and additional genes most closely related to AOA genomes (Fig. 4A). Another AOA virus contig (JGI virus accession no. RCM31_10001047), recovered from planktonic microbial communities in the Amazon River plume, also contained a *pcy* gene flanked by viral genes and most similar to those of AOA (Fig. 4A). In addition, two further viral contigs from saline lake (3300031223_assembled_Ga0307981_1000533) and marine (3300009371 assembled Ga0118717_1001322) environments encoded AOA-type *pcy* genes, both similarly flanked by viral genes (Fig. 4A). These consistent patterns strongly support the possibility of virus-mediated horizontal transfer of AOA pcy genes across distinct environments. Beyond the separate plastocyanin, plastocyanin can also be integral into AOA respiratory chain complexes III and IV, and fused in multicopper oxidase lineage 4 (MCO4) (Fig. 4A), although the physiological functions of MCO4 remain poorly understood ^25^. Phylogenetic analysis showed that plastocyanin proteins encoded by these four viral contigs clustered within a well-supported clade alongside plastocyanin sequences from terrestrial and marine AOA, clearly separated from other archaeal 1 and 2 domain cupredoxin-like proteins (Figs. 4C and D and Supplementary Fig. S13).

**Figure 4.**
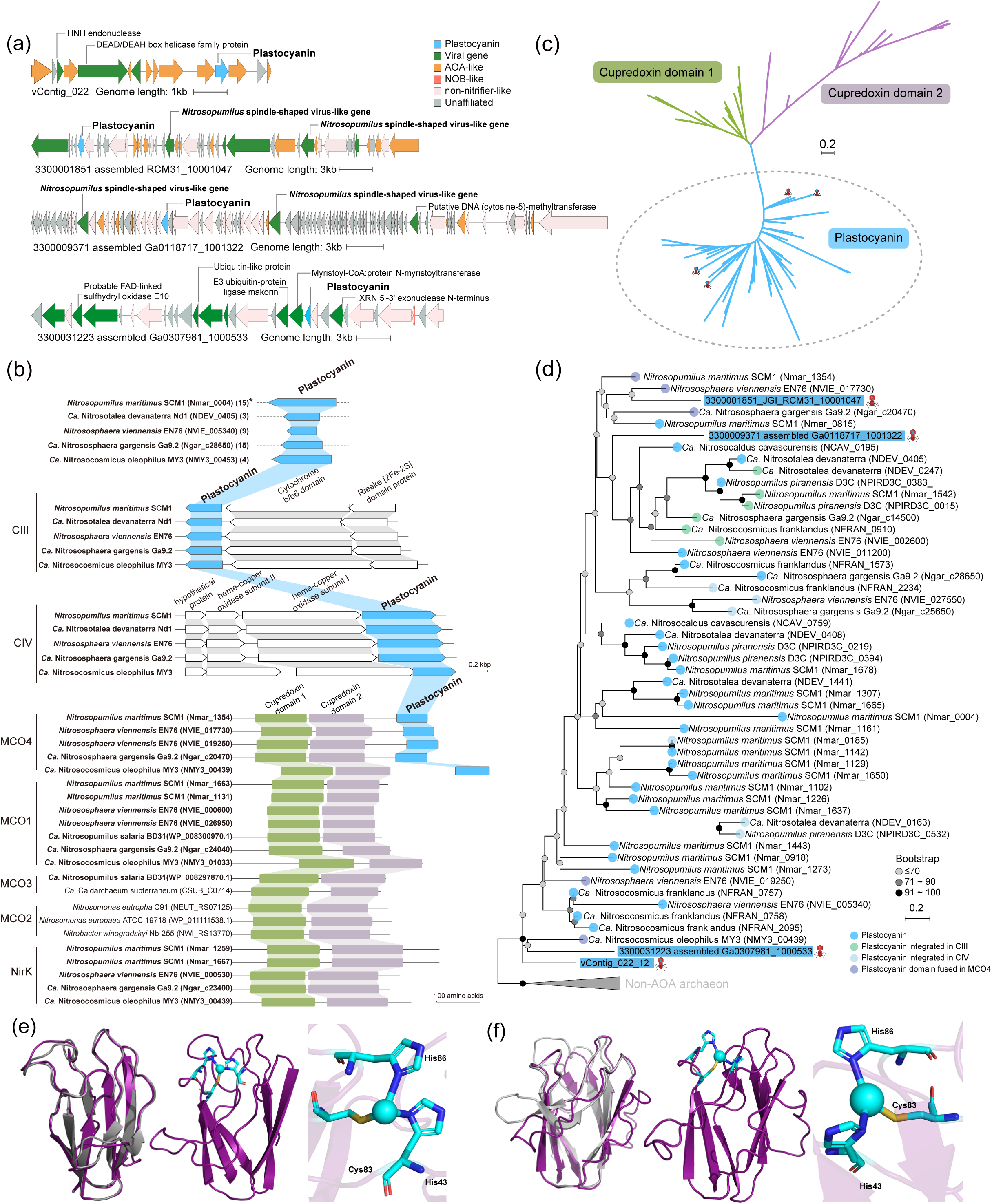
Genetic organization, phylogenetic relationship, and predicated 3D structures of viral plastocyanin proteins from AOA-infecting viruses. **(A)** Genetic maps of viral contigs predicated to originate from two AOA-infecting viruses (vContig_022 and RCM31_10001047) and two without host prediction (Ga0118717_1001322 and Ga0307981_1000533). *pcy* in three of them were flanked by viral genes. **(B)** Schematic presentation of plastocyanin proteins, plastocyanin and other cupredoxin domains integrated into multicopper oxidases (MCOs), NirK and electron transfer chain complexes III and IV in genomes of AOA and certain non-AOA prokaryotes. *Indicates the number of separate plastocyanin genes in each AOA genome, excluding plastocyanin domains integrated into MCO4 and complexes III and IV. **(C)** Simplified phylogeny of plastocyanins and cupredoxin domains integrated into MCOs, NirK, complex III or IV of the electron transfer chain (see Supplementary Fig. S12 for the full tree). **(D)** A maximum-likelihood phylogenetic tree of plastocyanin from AOA-infecting viruses and AOA genomes was built using IQ-TREE (version 2.1.2) ^84^ with the selected best-fit model (rtREV+I+G4). NCBI accession numbers are provided in parenthesis. Circles at nodes indicate bootstrap values from 1000 replicates, and the scale bar indicates 0.3 substitutes per position. **(E and F)** Predicted 3D structures of viral plastocyanin proteins from vContig_022 **(E)** and RCM31_10001047 from JGI IMG/VR v4 **(F)**. Middle images in the panel E and F present the predicted 3D structures of viral plastocyanin proteins modelled by I-TASSER-MTD (https://seq2fun.dcmb.med.umich.edu/I-TASSER/) ^85^ with default parameters. The left images illustrate the matches between the viral plastocyanin and best hit crystal structure of Nmar_1307 (grey, PDB accession code 5FC9) from the PDB library, with a Cov value 0.99 and an average template modeling score (TM-score) 0.92 for (E) and a Cov value 0.82 and a TM-score 0.74 for (F), as predicated by I-TASSER-MTD and visualized using PyMOL (version 3.12) ^86^. Right images in panel d and e zoom in on the copper-binding site, displaying the primary ligands as cyan sticks and the Cu ion as cyan sphere, consistent with the residue numbering in crystal structure of Nmar_1307.

The X-ray crystal structure of a plastocyanin protein from *N. maritimus* SCM1 (Nmar_1307; PDB accession code 5FC9) revealed a single copper ion coordinated in a trigonal plane by one cysteine (S_γ_Cys83-Cu = 2.25 Å) and two histidine residues (N_δ_His43-Cu = 2.01 Å and N_δ_His86-Cu = 2.07 Å) ^105^. These three copper-coordinating sites are conserved across all plastocyanin proteins encoded by both AOA-infecting viral contigs and AOA genomes (Supplementary Fig. S14). Predicted crystal structures of the two AOA-like plastocyanins from AOA-infecting viral contigs closely aligned with the characterized X-ray crystal structure of Nmar_1307 (with Cov value 0.99 and TM-score 0.92), revealing conserved copper-binding ligands typical of plastocyanins (Fig. 4E and F). Given that plastocyanins play a key role in AOA copper-dependent respiration, the acquisition of plastocyanin genes by viruses likely represents a strategy to modulate or augment host electron transport pathways, potentially benefiting both viral replication and host metabolic resilience under redox-variable conditions.

## Discussion

Global warming is driving an increase in the intensity, frequency and duration of heatwaves worldwide, with profound effects on microbial communities and activities, thereby altering biogeochemical cycles and ecosystem functions. Nitrifying microorganisms control the transformation of fixed nitrogen species and fix carbon via chemosynthetic pathways, with an important contribution to global nitrogen and carbon cycling. Interactions between nitrifiers and viruses, the most abundant biological entities on Earth, are emerging as key factors regulating nitrifier ecology and functioning ^44, 45, 48–54, 106^. However, how heatwaves influence nitrifier communities and their viral interactions across diverse environments remain poorly understood, reflecting a largely unexplored intersection of bottom-up (temperature-driven) and top-down (viral infection-mediated) controls on microbial ecology. We conducted incubation experiments under an extended heatwave condition of eight weeks, matching the maximum duration observed locally ^9, 13^. Our study revealed that heatwave temperature regimes substantially altered the composition of active members of nitrifying communities in tidal flats, with patterns distinct from those in adjacent agricultural soil ecosystems. The ^13^C-labeled nitrifier-infecting viruses exhibited temperature sensitive infection intensities, shifts in lytic and lysogenic lifestyles, and distinct AMG functional repertoires in coordination with their nitrifier hosts, highlighting the potential for heatwaves to reshape nitrifier-virus interactions and associated ecosystem functions. Notably, some AOA-infecting viruses carried *pcy* genes encoding key copper-dependent electron carriers involved in the AOA respiratory system, potentially modulating or supplementing electron transport under changing environmental conditions. Additionally, closely related viruses- and in some instances, even a single virus-were found to infect both AOB and NOB, which together drive two consecutive, metabolically interdependent steps of the nitrification process. These findings uncover a previously underappreciated layer of complexity in nitrifier-virus interactions under climate extremes, with important implications for understanding and predicting nitrogen and carbon cycling in land-sea transitional ecosystems in a warming world.

Coastal tidal flats are dynamic transition zones that harbor microbial communities originating from both adjacent marine and terrestrial ecosystems. These habitats host functionally redundant nitrifier ecotypes that coexist despite differing sensitives to major environmental variables such as temperature and salinity. This study revealed that sustained heatwave exposure triggered a pronounced shift of metabolically active nitrifying members in tidal flats, transitioning from typical marine ecotypes of AOA (NP-Gamma and NP-Theta), AOB (*N. marina*/*N. aestuarii*) and canonical NOB (*Nitrospira* lineage IV) to their respective terrestrial ecotypes (NS-Zeta and NS-Alph; *N. communis*/*N. nitrosa*.; and *Nitrolancea*). These terrestrial nitrifier ecotypes are known to exhibit greater thermal tolerance than their marine counterparts ^35, 36, 92, 96, 98, 99, 107^. Despite successfully occupying the niche following the replacement of marine nitrifiers under heatwave conditions, the limited salinity tolerance of these terrestrial ecotypes likely constrained their activity in the saline tidal flat environment ^92, 96, 98, 108, 109^, leading to reduced nitrification as indicated by lower NO_X_ accumulation. In contrast, in adjacent agricultural soils, exposure to sustained heatwave-like temperatures from 28°C to 37°C resulted in a clear shift in ^13^C-labeled active nitrifiers, with dominant soil AOB being replaced by terrestrial AOA lineages. These AOA ecotypes are known to encode thermotolerance-associated genes, and thus exhibit higher optimal growth temperatures than their terrestrial bacterial counterparts ^31, 35, 36^. Unlike in tidal flats, the absence of salinity stress in agricultural soils may have allowed these thermotolerant AOA ecotypes to thrive under elevated temperatures, leading to increased nitrification activity. These results highlight that land-use context plays a critical role in modulating the impact of heatwaves on nitrifying communities and their ecosystem functions.

Beyond nitrifier community shifts, our results indicate that heatwave conditions restructured virus-nitrifier host interactions at multiple functional levels. At 28°C, elevated virus-to-host ratios along with enrichment of lytic nitrifier infecting viruses and AMGs associated with energy metabolism, cofactors, and vitamin metabolism, and signaling proteins, likely reflected a virus-driven exploitation strategy aimed at maximizing replication through host metabolic takeover ^110, 111^. In contrast, the high temperature of heatwave (37°C) was associated with a decline in viral activity, a higher relative frequency of lysogenic phage derived from their lytic cycle, and the dominance of borderline AMGs related to the prokaryotic defense system, such as site-specific DNA-methyltransferase and type II toxin-antitoxin systems. These genes potentially offer the host additional protection against superinfection and aid in coping with environmental stress, and even in biofilm formation ^103^. Notably, the observed higher ratio of lysogeny to lytic viruses at 37°C does not necessarily reflect a lifestyle switch toward lysogeny under high temperature. An alternative explanation is that lysogeny was more frequently induced into the lytic cycle under heat stress, releasing free viral particles that were subsequently inactivated or degraded more rapidly in warmer conditions ^16, 112^, resulting in an apparent increase in the relative frequency of lysogeny. This interpretation is consistent with previous studies showing that higher temperatures are associated with reduced viral proliferation and increased inactivation rates ^113, 114^. Collectively, this coordinated shift between viruses and their nitrifier hosts underscores a temperature-dependent restructuring of virus-host dynamics, with significant implications for microbial community resilience and nutrient cycling in land-sea transitional ecosystems.

Copper plays a critical role in regulating AOA metabolism, with plastocyanin-encoding a type 1 blue copper-binding domain serving as a key electron carrier widely encoded across AOA lineages and essential for their copper-centric respiratory systems ^25, 115, 116^. In this study, we identified AOA-specific plastocyanin genes within viral contigs. Phylogenetic analysis and domain alignments showed that these viral plastocyanin sequences clustered closely with those from AOA genomes and shared conserved copper-binding sites, supporting a potential functional role in electron transfer. AOA typically harbor multiple copies of plastocyanin genes across lineages. For instance, the marine AOA model species *Nitrosopumilus maritimus* strain SCM1 encodes 17 small blue copper-containing plastocyanins, representing multiple homologs that are differentially expressed under varying growth conditions ^46^. This suggests functional diversification of plastocyanins, potentially enabling AOA to fine-tune their electron transport systems in response to environmental fluctuations. The presence of plastocyanin genes in AOA-infecting viruses may thus reflect a viral strategy to supplement or stabilize host electron transport, enhancing host adaptability and potentially promoting viral propagation. In addition to plastocyanins, Lee et al. recently reported that AOA-infecting viruses encode multicopper oxidase lineage 1 (MCO1) genes ^48, 49^. Analogous to cyanophages that carry AMGs encoding key photosystem components to boost host photosynthetic efficiency under stress ^117^, or to phages that harbor nitrogen fixation genes to induce heterocyst formation in *Anabaena* and *Nostoc* under nitrogen-limited conditions ^118^, AOA viruses may broadly transduce genes encoding copper-containing proteins involved in electron transfer and core energy metabolism in AOA for mutual benefit of both host and virus.

Ammonia oxidizers and nitrite oxidizers often coexist in close proximity or form aggregates in the environment to establish syntrophic associations ^119–122^, where AOB/AOA supply nitrite as an energy source for NOB, while NOB help alleviate nitrite toxicity for AOB/AOA or provide ammonia through urea hydrolysis ^123^.

Interestingly, in our study we now provide data that support the hypothesis that AOB and NOB share similar viruses, or even a single virus could infect multiple nitrifier hosts. This is consistent with the findings of recent studies that a single virus could infect microbial hosts that belong to phylogenetically distantly related microbial domains^124, 125^, especially those in aggregates or biofilms, such as the ANME (anaerobic methanotrophic archaea)-SRB (sulfate-reducing bacteria) or SOB (sulfur-oxidizing bacteria) -SRB pairs found in dense hydrothermal mats ^125^. Surveys of public datasets have also identified viruses capable of interacting with hosts across domains in diverse ecosystems characterized by syntrophic biofilms ^125^. Such patterns suggest that physical proximity and shared metabolic networks within aggregates may promote viral encounters with phylogenetically diverse hosts, eventually selecting for viruses with broader host ranges through ecological and evolutionary mechanisms, such as repeated co-infection and spatial constraints ^125^. Notably, these broad-host-range viruses may also facilitate lateral gene transfer (LGT) among co-occurring nitrifiers. For instance, the cyanase gene in the terrestrial AOA *Nitrososphaera gargensis* is phylogenetically closely related to those found in NOB ^27, 102^, implying that virus-mediated interdomain gene exchange cannot be excluded ^124, 125^. Palomo et al., based on reconciliation model, even suggested that an ancestor of *Nitrospira* (NOB) may have evolved into comammox *Nitrospira* by acquiring key ammonia oxidation genes, such as *amoA* and *haoA*, from β-proteobacterial AOB ^126^. Our study provides the first evidence that a single virus can potentially infect both AOB and NOB, offering new insights into virus-mediated interactions, gene flow, and coevolution within syntrophic nitrifying communities.

In conclusion, our study highlights the profound impact of prolonged heatwaves on the restructuring of active nitrifying communities and their interactions with viruses in tidal flats and agricultural soils, with direct consequence for the nitrification process in these ecosystems. Our findings suggest that sustained heatwaves substantially reduce the resistance of nitrifier communities, raising the key question of whether these communities can exhibit resilience by recovering to their original state once the disturbance subsides. While heatwaves can occur independently, they sometimes cooccur with drought, referred to as dry heatwaves, or with high humidity, especially in coastal regions, leading to so-called humid heatwaves ^127^. Further research is needed to elucidate the combined effects of heatwave with drought or humid conditions on nitrogen-cycling microbial communities and their functional outputs. Moreover, beyond nitrification, heatwaves also impact microbial processes linked to other nutrient cycles, greenhouse gas emissions, and carbon sequestration ^18^, thereby contributing to biogeochemical feedback loops that may further amplify climate change. Therefore, advancing our understanding of these mechanisms will improve the accuracy of models and projection regarding the interactions between extreme climate events and ecosystem services, which are ultimately underpinned by the resistance, resilience, and functionality of microbial communities.

## Acknowledgements

This work was supported by National Natural Science Foundation of China grants 42477318 and 42277304 (to B.W.), and U22A20590 (to Y.G.). This work was also funded by the US Department of Energy Early Career Research Program (DE-SC0025455 to W.Q.) and the startup funds of the University of Illinois Urbana-Champaign and the University of Oklahoma to W.Q.

## Notes

### Competing Interest Statement

The authors have declared no competing interest.

